# Inhibition of Centrosome Clustering Reduces Cystogenesis and Improves Kidney Function in Autosomal Dominant Polycystic Kidney Disease

**DOI:** 10.1101/2022.11.16.516801

**Authors:** Tao Cheng, Aruljothi Mariappan, Ewa Langner, Kyuhwan Shim, Jay Gopalakrishnan, Moe R. Mahjoub

## Abstract

Autosomal Dominant Polycystic Kidney Disease (ADPKD) is an inherited monogenic disorder accounting for ∼5% of patients with renal failure. Yet, therapeutics for the treatment of ADPKD remain limited. ADPKD tissues display defects in the biogenesis of the centrosome which causes genome instability, aberrant ciliary signaling, and secretion of pro-inflammatory factors that drive cyst growth and fibrosis. Cystic cells form excess centrosomes via a process termed centrosome amplification (CA), which often causes abnormal multipolar spindle configurations, mitotic catastrophe, and reduced cell viability. However, cells with CA can suppress multipolarity via “centrosome clustering”, a key mechanism by which cells circumvent apoptosis. Here, we demonstrate that inhibiting centrosome clustering can counteract the proliferation of renal cystic cells with high incidences of CA. Using ADPKD human cells and mouse models, we show that blocking centrosome clustering with two inhibitors, CCB02 and PJ34, blocks cyst initiation and growth *in vitro* and *in vivo*. Inhibition of centrosome clustering activates a p53-mediated mitotic surveillance mechanism leading to apoptosis, reduced cyst expansion, interstitial fibrosis, and improved kidney function. Transcriptional analysis of kidneys from treated mice identified pro-inflammatory signaling pathways implicated in CA-mediated cystogenesis and fibrosis. Our results provide the first evidence that centrosome clustering is a cyst-selective target for the improvement of renal morphology and function in ADPKD.

## Introduction

Autosomal Dominant Polycystic Kidney Disease is the most common inherited monogenic kidney disorder, affecting ∼13 million individuals worldwide. ADPKD is characterized by dysregulation of renal epithelial cell homeostasis and hyperproliferation of normally quiescent cells, which profoundly alters organ architecture and impairs renal function. It is well established that ADPKD is a ciliopathy, a disease caused by ciliary dysfunction. Cilia are microtubule-based organelles that play important chemo- and mechanosensory roles in cells (*1*). In the kidney, cilia are found on renal progenitor cells during development (*2, 3*), and quiescent epithelial cells lining the various segments of mature nephrons (*4*). The cilia protrude from the apical surface and are in contact with the extracellular environment, acting as a cellular sensor that regulates epithelial cell growth, homeostasis and repair (*5, 6*). The assembly of cilia is templated by the centrosome, the main microtubule-organizing center in animal cells (*7*). Most cells in the human body, including in the kidney, contain a solitary centrosome and cilium, and cells have evolved tight regulatory mechanisms to ensure they contain only one of each organelle (*8*).

Recent studies have noted the presence of excess centrosomes (a phenomenon termed **C**entrosome **A**mplification; CA) in renal epithelial cells of ADPKD patients (*9, 10*). CA is most commonly caused by centrosome overduplication, due to uncoupling of the centrosome biogenesis process from the cell cycle (*8*), and is detrimental to cell physiology in multiple ways: (a) it causes abnormal mitotic spindle formation leading to chromosome segregation errors and genome instability (*11, 12*), (b) disrupts cilia assembly and signaling (*13*), and (c) results in increased secretion of cytokines and pro-inflammatory paracrine signaling factors (*14, 15*). Collectively, these changes enhance the proliferative capacity of cells (and tissues) associated with CA. This phenomenon has been extensively studied in the context of cancer, as CA has been shown to play a critical role in driving tumor cell proliferation and metastasis (*15*). Intriguingly, ADPKD is also caused in part by defects in genome stability, ciliary function, and pro-inflammatory signaling (*16*). Yet it has remained unclear whether the CA observed in kidney samples of ADPKD patients contributes to renal cystogenesis, or how it affects kidney physiology.

To address these questions, we recently examined the consequences of CA during kidney development, homeostasis and repair post-injury. Analysis of kidney specimens from patients with ADPKD showed that CA is highly prevalent in cystic epithelial cells but extremely rare in healthy kidneys (*17*). CA was evident early during cyst formation, and even present in pre-cystic tubules, indicating this defect likely precedes cystogenesis. Using a genetic mouse model with which we can alter centrosome biogenesis in kidneys *in vivo*, we demonstrated that the formation of excess centrosomes indeed caused rapid-onset cystogenesis (*17*). Genetic induction of CA during embryonic kidney development in wild-type mice caused defects in the growth and differentiation of renal progenitors, resulting in kidneys that were smaller than normal at birth. Importantly, the kidneys developed cysts very rapidly, resulting in kidney failure and lethality by postnatal day 15. Analysis of cells with CA showed defects in mitotic spindle morphology, ciliary assembly, and signaling pathways essential for growth and differentiation of renal progenitors, highlighting the mechanisms underlying the observed developmental phenotypes (*17*). Moreover, CA sensitized kidneys in adult mice, causing extensive cyst formation within 30 days after ischemic renal injury (*17*). Thus, the formation of excess centrosomes can be a driver of renal cyst formation during kidney development and homeostasis.

In addition to the cell-intrinsic defects (i.e. genome instability and ciliogenesis) that confer a proliferative advantage to themselves, cells with amplified centrosomes have been found to secrete high levels of cytokines and growth factors, resulting in pro-inflammatory and proliferative paracrine signaling (*14, 15*). In essence, cells with CA can act as “signal amplifiers” that promote the growth of adjacent cells with normal centrosome number (*14, 15*). Thus, it has been proposed that eliminating cells with CA will attenuate both cell intrinsic and extrinsic effects. Recent studies in cancer cells support this theory. Amplified centrosomes typically cause the formation of multipolar mitosis and chromosome mis-segregation errors, leading cells to undergo mitotic catastrophe and cell death (Figure 1A). However, cancer cells with extra centrosomes achieve a pseudo-bipolar spindle configuration via a process termed “centrosome clustering”, a key adaptive mechanism by which they circumvent mitotic catastrophe and survive. Thus, inhibiting centrosome clustering to induce multipolar divisions, leading to negative selection of that population of cells, has been proposed as a strategy to counteract tumors with high incidences of centrosome amplification (Figure 1A) (*18*). To achieve this a number of drug discovery screens have identified selective small molecule inhibitors of centrosome clustering, some of which are currently in clinical trials for different types of human cancers (*19, 20*).

**Figure 1.**
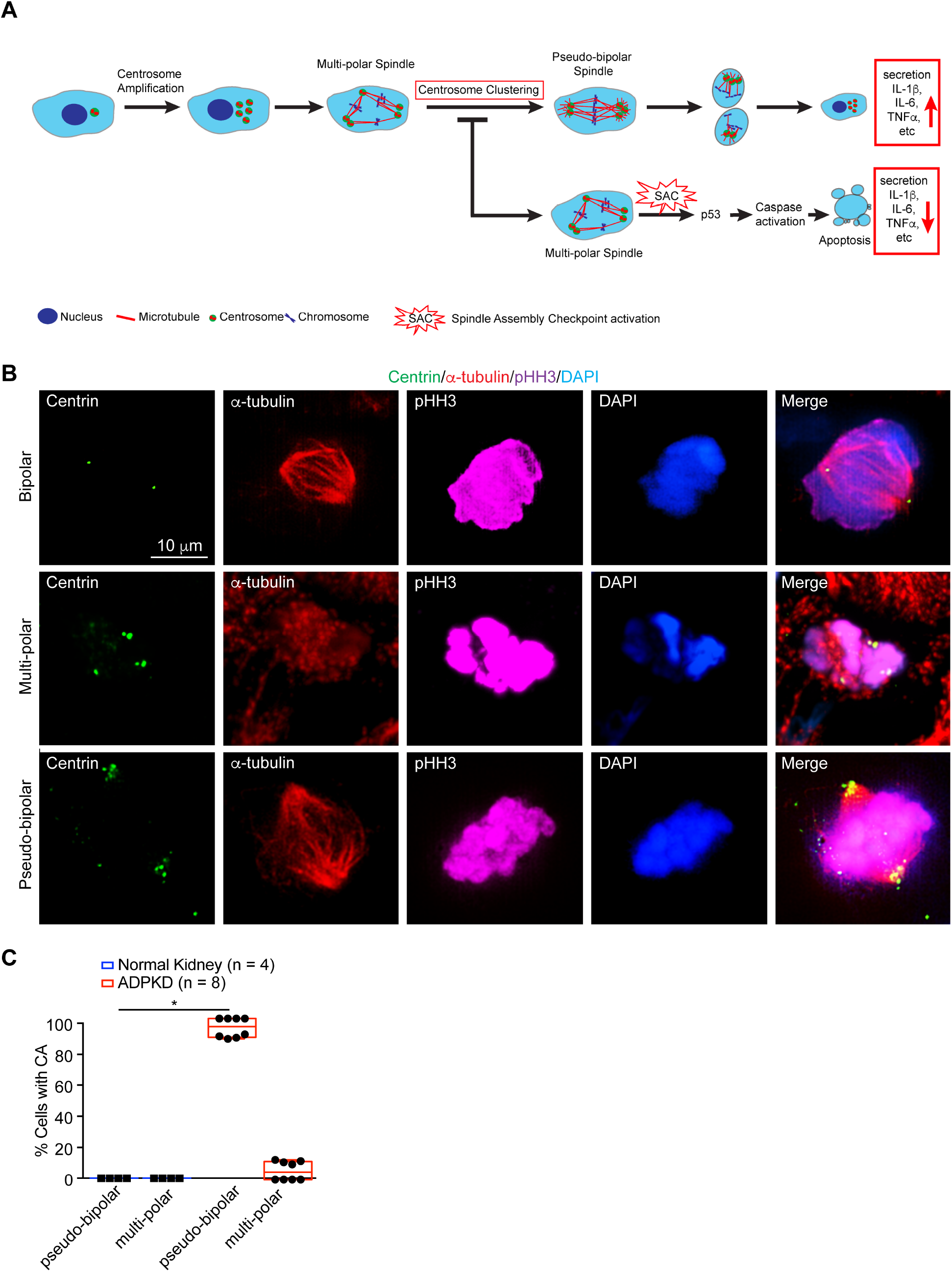
Human ADPKD cells with amplified centrosomes form pseudo-bipolar mitotic spindles. **(A)** Cartoon schematic of centrosome amplification and its consequences. Mitotic spindles in cells with excess centrosomes typically form more than two poles, leading to multipolar spindle configurations and activation of the Spindle Assembly Checkpoint. This is followed by activation of the p53 mitotic surveillance pathway, leading to caspase-mediated apoptosis. Centrosome clustering and the formation of pseudo-bipolar spindles is a survival mechanism adapted by some of these cells to avoid cell death. These cells subsequently demonstrate enhanced secretion of pro-inflammatory cytokines and growth factors. Inhibition of centrosome clustering can push cells towards the apoptotic pathway. **(B)** Immunofluorescence staining of human ADPKD kidney sections with antibodies to highlight centrioles (centrin), spindle microtubules (α-tubulin) and DNA (phospho-Histone H3 and DAPI). **(C)** Quantification of spindle configurations in dividing cells from wild-type and ADPKD kidneys. * = p<0.05 (one-way ANOVA).

In this study, we sought to determine whether pharmacological inhibition of centrosome clustering in ADPKD mice can result in elimination of cells with CA, attenuation of progressive cystic growth and improvement in kidney function *in vivo*.

## Results

### ADPKD cells with amplified centrosomes form pseudo-bipolar mitotic spindles.\

We previously demonstrated that CA is prevalent in cyst-lining epithelia of ADPKD patient kidneys (*17*). Roughly 12% of cystic epithelial cells contain excess centrosomes (ranging from 5-23%), and this phenotype is evident in ∼43% of cysts (*17*). This range is consistent with what has been observed in numerous types of cancer cells with CA (*21-25*). To determine whether dividing cystic epithelia with CA clustered their centrosomes in mitosis, ADPKD patient kidney samples were immunostained with antibodies targeting centrosomes, microtubules and the mitotic marker phospho-Histone H3. Mitotic cells with normal centrosome number formed bipolar spindles as expected, whereas cells with CA almost exclusively formed pseudo-bipolar spindle configurations in both metaphase and anaphase (Figure 1, B and C). This indicates that a robust centrosome clustering mechanism exists in ADPKD cystic epithelia, likely contributing to the ability of these cells to survive mitotic catastrophe, proliferate and form cysts.

### Inducing centrosome amplification accelerates the cystic disease phenotype in a slow-onset mouse model of ADPKD

We previously demonstrated that inducing the formation of excess centrosomes in wild-type mice caused rapid cystogenesis shortly after birth (between P0-P15) (*17*). To determine the synergistic effects of CA in conjunction with loss-of-function in ADPKD genes, we utilized a recently developed slow-onset ADPKD mouse model harboring a point mutation in *Pkd1* (*26*). The *Pkd1* variant p.Arg3277Cys (RC) in humans is hypomorphic and associated with typical slow-onset ADPKD in homozygosity (*26*). A mouse mimicking this variant (hereafter referred to as *Pkd1*^*RC/RC*^) is viable for at least one year with slowly progressive PKD. Cysts develop spontaneously, kidneys enlarge slowly up to 6 months of age and cystogenesis accelerates beginning at 9 months. Interstitial renal fibrosis - a key factor that causes of scarring and renal failure in ADPKD - and decline in filtration function start at 6 months of age and similarly accelerate at 9 months (*26, 27*). Quantification of centrosome number in 5-month-old *Pkd1*^*RC/RC*^ mice shows that between 5-15% of cystic epithelial cells contain excess centrosomes (Figure 2D and Supplemental Figure 1C), consistent with what we observed in human ADPKD samples. Overall, this mouse model effectively mimics the pathophysiological features of slow–onset progressive cystogenesis of human ADPKD, provides a large window of time to study the disease progression, and has been used extensively for preclinical testing of potential therapeutic compounds (*26, 28-42*).

**Figure 2.**
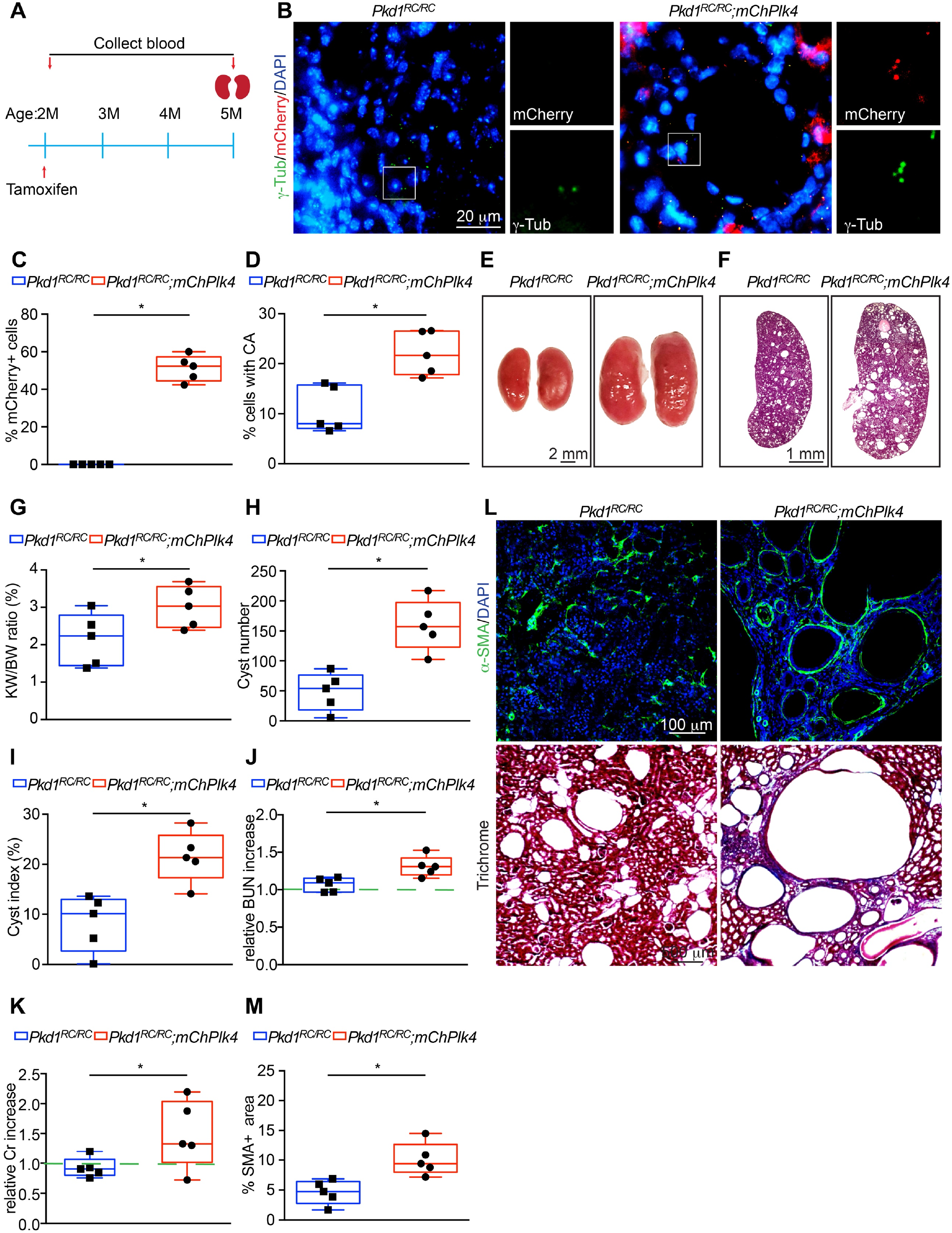
Induction of centrosome amplification accelerates cystogenesis in slow onset ADPKD mice. **(A)** Timeline of conditional induction of centrosome amplification in *Pkd1*^*RC/RC*^*;mChPlk4* mice, including serum and kidney sample collection. **(B)** Immunofluorescence staining of kidney sections from 5-month-old *Pkd1*^*RC/RC*^ and *Pkd1*^*RC/RC*^*;mChPlk4* mice using antibodies against mCherry, centrosomes (*γ*-tubulin) and DNA (DAPI). **(C)** Quantification of the percentage of mChPlk4-positive cells and **(D)** the percentage of cyst-lining cells with excess (> 2) centrosomes. **(E** and **F)** Images of whole kidneys and H&E-stained sections of *Pkd1*^*RC/RC*^ and *Pkd1*^*RC/RC*^*;mChPlk4* mice at 5 months of age. **(G)** Evaluation of kidney weight expressed as percentage of body weight at 5 months. **(H)** Quantification of cyst number and **(I)** fractional cyst area per kidney section. **(J)** Analysis of relative blood urea nitrogen (BUN) and **(K)** serum creatinine levels at 5 months. **(L)** Immunofluorescence staining (top) with α-smooth muscle actin (SMA; to mark myofibroblasts) and DNA (DAPI), and Trichrome staining (bottom) of kidney sections from 5-month-old *Pkd1*^*RC/RC*^ and *Pkd1*^*RC/RC*^*;mChPlk4* mice. (**M)** Quantification of the fraction of α-SMA-positive area. N = 5 mice per group for all experiments. * = p<0.05 (one-way ANOVA).

To cause elevated levels of centrosome amplification in the ADPKD mutant background, *Pkd1*^*RC/RC*^ were crossed with a transgenic model that allows for conditional expression of mCherry-tagged Plk4, which we previously used to drive CA in wild-type mouse kidneys (*17*). Expression of Plk4 was induced in *Pkd1*^*RC/RC*^;*mChPlk4* animals beginning at 2 months of age (Figure 2A) when cystogenesis is mild, the kidney weight-to-body weight (KW/BW) ratio is normal, and kidney function is mostly unaffected (*26*). This provided a means to examine the effects of enhanced CA at stages when the ADPKD phenotype is mild in this model. Immunofluorescence staining of kidneys isolated after 3 months confirmed overexpression of Plk4, and a concurrent 55% increase in the fraction of cells with CA (Figure 2, B and D) compared to *Pkd1*^*RC/RC*^ mice alone. The kidneys became larger in size, with a ∼35% increase in KW/BW (Figure 2, E and G, and Supplemental Figure 1A). Quantification of cyst number and fractional cyst area showed a near 175% and 120% increase, respectively (Figure 2, F, H and I and Supplemental Figure 1B). Analysis of blood urea nitrogen (BUN) and serum creatinine indicated elevated levels compared to the baseline in *Pkd1*^*RC/RC*^ mice, highlighting a decline in kidney filtration function upon CA (Figure 2, J and K, Supplemental Figure 1, D and E). This was accompanied by a ∼102% increase in interstitial fibrosis (Figure 2, L and M). Together, these data indicate that centrosome amplification can significantly accelerate the cystic disease phenotype in slow-onset ADPKD mice.

### Inhibition of centrosome clustering in ADPKD cells activates the SAC and blocks proliferation

Next, we sought to test whether blocking centrosome clustering in ADPKD cells with excess centrosomes can halt their proliferation, as has been proposed for cancer cells. Three inhibitors of centrosome clustering that have recently been identified in high-throughput *in vitro* screens - and also tested in mice - are CCB02, AZ82 and PJ34. CCB02 is a novel tubulin-binding molecule that competes for the tubulin-binding site of the centrosomal protein CPAP, forcing the extra centrosomes in a cell to nucleate high levels of microtubules prior to mitosis and thus prevent them from clustering (*43*). Treatment of various cancer cells with CCB02 reduced the fraction with supernumerary centrosomes *in vitro* and resulted in decreased cell proliferation. Notably, CCB02 had no detrimental effect on dividing cells with the normal complement of centrosomes (*43*). Oral administration of CCB02 to mice bearing human cancer xenografts resulted in reduction of centrosome amplified tumor cells *in vivo*. Blocking centrosome clustering caused significant inhibition of tumor growth at doses well tolerated by the animals (*43*). AZ82 is an inhibitor of KIFC1, a kinesin-14 family motor protein that has been shown to play a critical role in centrosome clustering in various types of cancer cells (*44, 45*). PJ34 is a phenanthrene-derived PARP1 inhibitor that can suppress the expression and activity of the kinesin KIFC1 (*46-48*). PJ34 treatment blocks centrosome clustering, induces multipolar spindle formation in cells with CA (*46*), and results in decreased breast cancer cell proliferation. Collectively, these findings indicate that blocking centrosome clustering with these inhibitors is a viable approach to reduce the proliferation of cells with CA *in vivo*.

Using these three compounds we first sought to test whether inhibition of centrosome clustering can induce the formation of multipolar mitotic spindles and mitotic arrest in ADPKD cells *in vitro*. Quantification of centrosome number in wild-type human kidney cells (HK-2) and *PKD1*-null cells (WT9-12) (*49, 50*) showed that roughly a third of ADPKD cells contained excess centrosomes, significantly higher than the control cells (Figure 3, A-C). As *PKD1*-null cells with CA enter mitosis, the vast majority (>80%) clustered their centrosomes and displayed a pseudo-bipolar spindle configuration (Figure 3, D and E). To assess how mitotic cells with excess centrosomes respond to the centrosome clustering inhibitors, both control and *PKD1*-null cells were treated with either CCB02, PJ34 or AZ82 for 24 h (Figure 3D). Consistent with previous reports showing no adverse effects on spindle formation in cells with normal centrosome number (*43-45, 47, 48*), both control and *PKD1*-null cells with two centrosomes predominantly formed bipolar spindles in metaphase when treated with the inhibitors (Figure 3, D and E). In contrast, *PKD1*-null cells with CA formed multipolar and disorganized spindles in mitosis (Figure 3, D and E). This was evident upon treatment with CCB02 and PJ34, but not significantly with AZ82. Since the AZ82 inhibitor showed cellular toxicity and caused lethality when tested in ADPKD mice (data not shown), we focused solely on CCB02 and PJ34 for the remainder of this study. In CCB02 and PJ34 treated cells, centrosomes were present at the majority of excess spindle poles and there was a concurrent decrease in the proportion of metaphase cells with bipolar spindles (Figure 3E).

**Figure 3.**
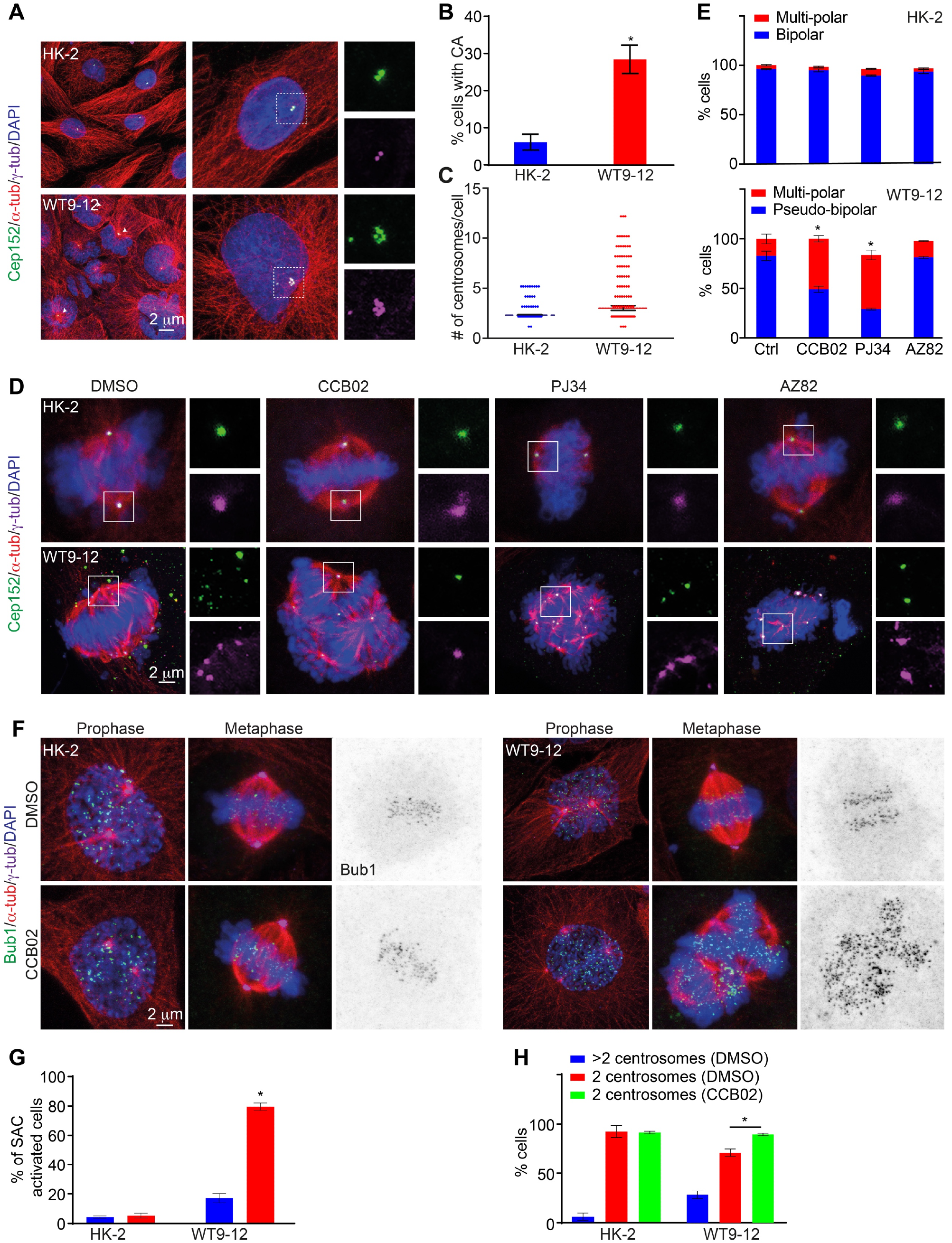
Inhibition of centrosome clustering in ADPKD cells promotes multipolar spindle formation in mitosis and activates the spindle assembly checkpoint. **(A)** Immunofluorescence staining of interphase wild-type (HK-2) and *pkd1*-null (WT9-12) cells with antibodies that mark centrioles (Cep152), centrosomes (γ-tubulin), microtubules (α-tubulin) and DNA (DAPI). **(B)** Quantification of the percentage of cells with amplified centrosomes. N = 293 (HK-2) and 374 (WT9-12) cells. **(C)** Distribution of the number of excess centrosomes per cell. N = 301 (HK-2) and 322 (WT9-12) cells. **(D)** Immunofluorescence staining of mitotic wild-type (HK-2) and *pkd1*-null (WT9-12) cells with antibodies that mark centrioles (Cep152), centrosomes (γ-tubulin), microtubules (α-tubulin) and DNA (DAPI). Cells were treated with vehicle only (DMSO), CCB02, PJ34 or AZ82 for 24 h. **(E)** Top graph shows quantification of the percentage of wild-type cells that form bipolar (cells containing normal centrosome number) or multipolar (cells with >2 centrosomes) spindles. Bottom graph shows quantification of the percentage of *pkd1*-null cells with amplified centrosomes that formed pseudo-bipolar (clustered centrosomes) versus multipolar (de-clustered centrosomes). For HK-2 cells: N = 207 (control), 674 (CCB02), 174 (PJ34), 101 (AZ82); for WT9-12 cells: N = 393 (control), 449 (CCB02), 205 (PJ34), 189 (AZ82). **(F)** Immunofluorescence images of mitotic wild-type and *pkd1*-null cells stained with antibodies that mark centrosomes (γ-tubulin), microtubules (α-tubulin), Bub1 and DNA (DAPI). Cells were incubated with DMSO or CCB02 for 24 h. **(G)** Quantification of the percentage of cells showing Bub1 accumulation in mitosis. For HK-2 cells: N = 167 (DMSO), 133 (CCB02); for WT9-12 cells: N = 180 (DMSO), 278 (CCB02). **(H)** Quantification of the percentage of wild-type and *pkd1*-null cells containing normal (2) or excess (>2) centrosomes following treatment with CCB02. N = 371 (HK-2) and 264 (WT9-12) cells. All results are from three independent experiments. * = p<0.05 (two-way ANOVA).

Next, we tested whether CCB02-induced centrosome de-clustering caused a mitotic delay, activation of the spindle assembly checkpoint (SAC) and elimination of cells with CA from the population. Control and *PKD1*-null cells were incubated for 24 h with either vehicle or CCB02 and cells stained using antibodies against Bub1, which accumulates on unattached kinetochores and acts as a surrogate marker of SAC activation (*43, 51*). First, we confirmed that cells with either normal or excess centrosomes showed an accumulation of Bub1 in prophase (Figure 3F) which is expected due to the unattached kinetochores at this stage of the cell cycle (*52*). Cells with bipolar and pseudo-bipolar spindles showed a decrease in these levels in metaphase (Figure 3, F and G). In contrast, CCB02-treated *PKD1*-null cells with multipolar spindles displayed significant accumulation of Bub1 in metaphase (Figure 3, F and G), indicating that inhibition of centrosome clustering activates the SAC specifically in extra centrosome-containing cells. Finally, quantification of the fraction of cells with CA after incubation with centrosome clustering inhibitors showed a decrease in the percentage with greater than 2 centrosomes, and a concurrent increase in cells with the normal complement (Figure 3H). Collectively, these data indicate that inhibiting centrosome clustering in ADPKD cells promotes formation of multipolar mitotic spindles, activation of the SAC, and reduced abundance of cells with CA due to apoptosis.

### Inhibition of centrosome clustering attenuates cystic disease progression in vivo

To determine the effects of centrosome clustering inhibitors on cyst growth *in vivo, Pkd1*^*RC/RC*^ mice were first treated with CCB02 or PJ34 starting at 9 months of age, the stage when expansion of cystic area, cyst number and KW/BW begin to accelerate (*26*). Mice were administered CCB02 or PJ34 every two days for a total period of 2 months, which was well tolerated by the animals (Figure 4, A and B). Immunofluorescence analysis of kidney sections from control (vehicle only) *Pkd1*^*RC/RC*^ mice indicated that the majority (∼95%) of cells with CA displayed a clustered configuration (Figure 4, C and D). In contrast, both CCB02 and PJ34-treated kidney cells with CA predominantly contained multipolar spindles (Figure 4, C and D), indicating that the centrosome clustering inhibitors are also active *in vivo*. Remarkably, treatment with either inhibitor resulted in a significant decrease in kidney size, cyst index and cyst number compared to control (Figure 4E-I, Supplemental Figure 2,3). Quantification of BUN and serum creatinine showed elevated levels relative to the baseline in control *Pkd1*^*RC/RC*^ mice over that 2-month period, whereas mice treated with either inhibitor maintained significantly lower levels of both (Figure 4, J and K, Supplemental Figure 4, A and B). In addition, there was a concurrent decrease in interstitial fibrosis (Figure 4, L and M). These data indicate that inhibiting centrosome clustering during the rapid cyst-expansion stage results in reduced cyst growth, improved kidney morphology and preserved renal function.

**Figure 4.**
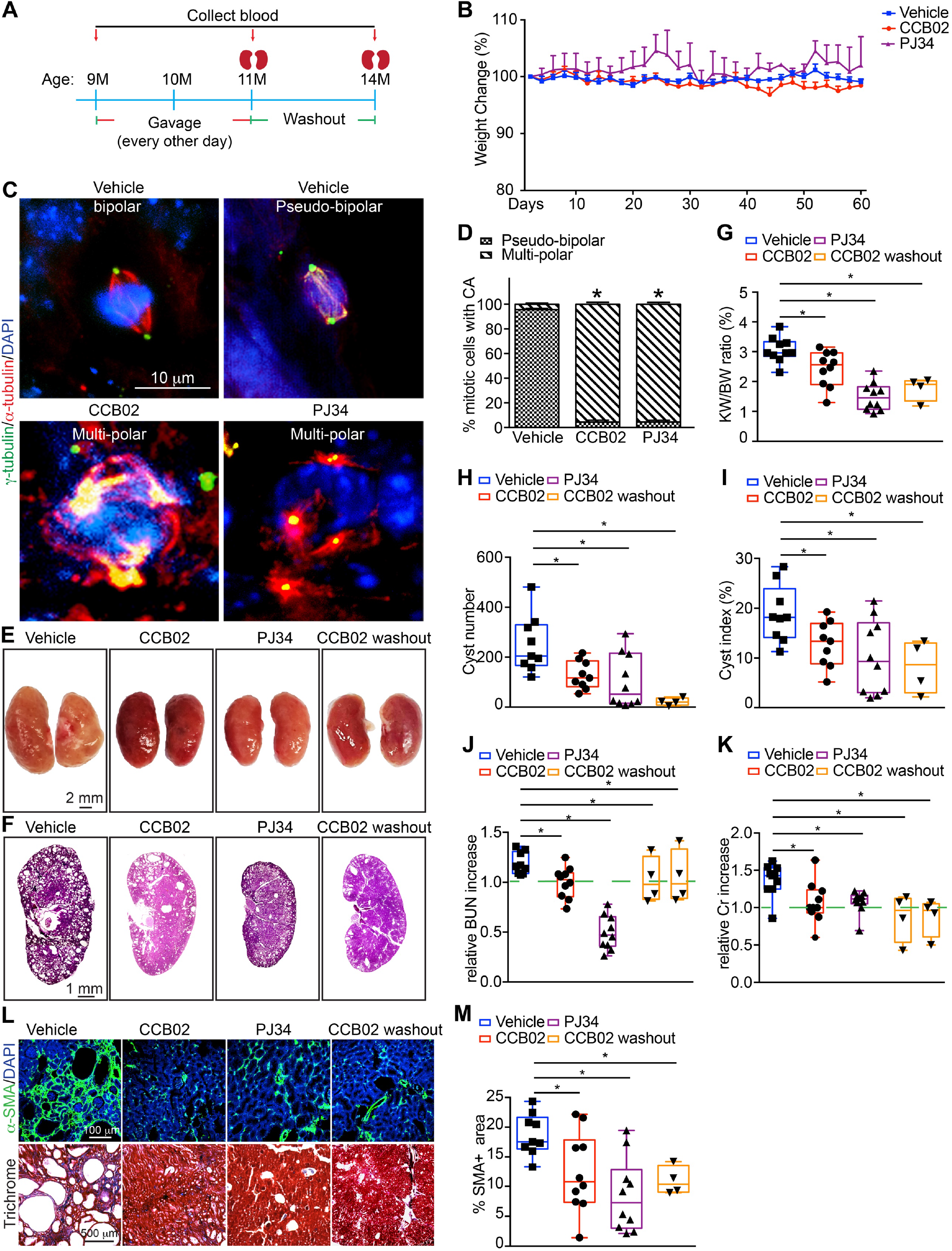
Inhibition of centrosome clustering attenuates disease progression during rapid stages of cytogenesis *in vivo*. **(A)** Schematic representation of CCB02 and PJ34 treatment scheme and washout in *Pkd1*^*RC/RC*^ mice. **(B)** Quantification of relative weight change during the treatment timeline. **(C)** Immunofluorescence staining of kidney sections from *Pkd1*^*RC/RC*^ mice at 11 months with antibodies to highlight centrosomes (γ-tubulin), spindle microtubules (α-tubulin) and DNA (DAPI). **(D)** Quantification of the percentage of spindle configurations at metaphase. N = 201 cells (vehicle group), 134 (CCB02 group), and 117 (PJ34 group). **(E** and **F)** Images of whole kidneys and H&E-stained sections of *Pkd1*^*RC/RC*^ mice at 11 months of age after treatment with CCB02 or PJ34, and post-washout (14 months). **(G)** Evaluation of kidney weight expressed as percentage of body weight. **(H)** Quantification of cyst number and **(I)** fractional cyst area per kidney section in treated and untreated mice. **(J)** Analysis of relative blood urea nitrogen (BUN) and **(K)** serum creatinine levels at 11 months and post-washout (14 months). **(L)** Immunofluorescence staining (top) with α-smooth muscle actin (SMA; myofibroblasts) and DNA (DAPI), and Trichrome staining (bottom) of kidney sections following the indicated treatment regimen. **(M)** Quantification of the fraction of α-SMA-positive area. N = 10 mice each for Vehicle, CCB02 and PJ34; N = 4 animals for CCB02 washout group. * = p<0.05 (one-way ANOVA).

Next, we sought to determine whether treatment with centrosome clustering inhibitors beginning earlier, and for a longer period of time, would yield better outcomes with regards to cystogenesis and kidney function. *Pkd1*^*RC/RC*^ mice at 6 months of age display relatively mild ADPKD phenotypes, with minimal increase in kidney volume, cyst size and number, nor a significant decline in renal function (*26, 27, 36*). Using a similar dosing scheme, mice were administered either vehicle or CCB02 beginning at 6 months of age every 2 days for a total of 5 months (Figure 5A). In parallel, *Pkd1*^*RC/RC*^ mice were given Tolvaptan, a selective vasopressin receptor 2 (V2) antagonist that is currently the only FDA approved drug for treatment of ADPKD (*53, 54*) and has a different mechanism of action than centrosome clustering inhibitors. Similar to the 2-month treatment plan, the mice showed no adverse effects to CCB02, whereas Tolvaptan-treated animals showed significant weight loss over that 5-month period (Figure 5B), which has been previously reported as a significant side effect of this compound (*54, 55*). Treatment with CCB02 resulted in preservation of kidney size, reduced cyst index and cyst number compared to control *Pkd1*^*RC/RC*^, and better than the Tolvaptan treated group (Figure 5C-G and Supplemental Figure 5, A and B). Importantly, these differences were more pronounced when compared to the 2-month treatment plan (Figure 4E-I, Supplemental Figure 2,3). Quantification of BUN, serum creatinine and interstitial fibrosis similarly showed improved preservation of kidney function over the 5-month period (Figure 5H-K, Supplemental Figure 5, C and D) when compared to the 2-month treatment plan (Figure 4J-M, Supplemental Figure 4, A and B). The maintenance of kidney function upon CCB02 treatment was similar to the effect of Tolvaptan, highlighting the potential for centrosome clustering inhibitors in reducing disease development and severity over prolonged periods.

**Figure 5.**
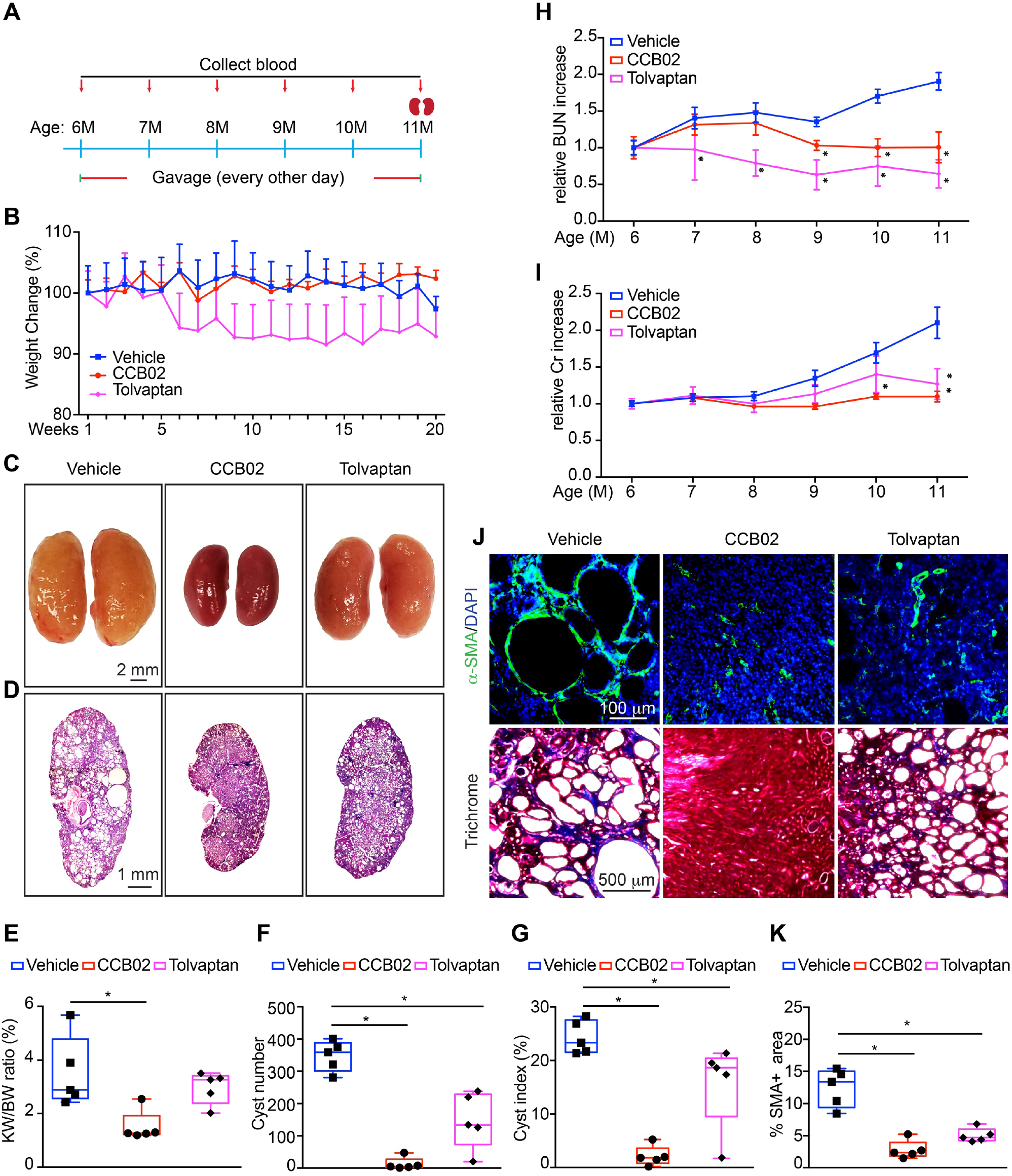
Inhibition of centrosome clustering at earlier stages of cystogenesis shows stronger effects on disease progression. **(A)** Schematic representation of CCB02 and Tolvaptan treatment scheme in *Pkd1*^*RC/RC*^ mice. **(B)** Quantification of relative weight change during the treatment timeline. **(C** and **D)** Images of whole kidneys and H&E-stained sections of *Pkd1*^*RC/RC*^ mice at 11 months of age after treatment with CCB02 or Tolvaptan. **(E)** Evaluation of kidney weight expressed as percentage of body weight. **(F** and **G**) Quantification of cyst number and fractional cyst area per kidney section in treated and untreated mice. **(H)** Analysis of relative changes in blood urea nitrogen (BUN) and **(I)** serum creatinine levels. **(J)** Immunofluorescence staining (top) with α-SMA (myofibroblasts) and DNA (DAPI), and Trichrome staining (bottom) of kidney sections following the indicated treatment regimen. **(K)** Quantification of the fraction of α-SMA-positive area. N = 5 mice per group for all experiments. * = p<0.05 (one-way ANOVA).

Finally, we sought to determine whether short-term treatment of ADPKD mice with centrosome clustering inhibitors, followed by a period of no drug administration, would provide beneficial long-term effects. Nine-month-old *Pkd1*^*RC/RC*^ mice were given CCB02 every other day for 60 days, the treatment was stopped, and the mice were followed for 3 months until they reached 14 months of age (Figure 4A). Analysis of kidney morphology, function and fibrosis showed that the dramatic improvements observed at 11 months were maintained, even when the drug was no longer administered (Figure 4E-M). In sum, our results indicate that inhibiting centrosome clustering in slow-onset ADPKD mice results in reduced cyst growth, improved kidney morphology and preserved renal function, which are evident even after drug treatment is halted.

### Centrosome clustering inhibitors promote apoptosis of cells with CA and reduced inflammatory signaling

Spindle formation defects caused by CA often result in activation of mitotic stress and DNA-damage sensors, p53-mediated cell cycle arrest, and ultimately cell death via induction of apoptotic signaling pathways (*17, 43, 56-59*). To determine whether inhibition of centrosome clustering promotes p53-mediated cell death, kidneys of *Pkd1*^*RC/RC*^ mice were immunostained for p53. We noted significant nuclear accumulation of p53 in mice treated with CCB02 or PJ34 for both 2- and 5-months, compared to control untreated animals (Figure 6, A and B and Supplemental Figure 6, A and B). This corresponded with an increase in cell death as noted by TUNEL staining (Figure 6, C and D and Supplemental Figure 6, C and D), and a concomitant reduction in the number of cells with amplified centrosomes (Figure 6, E and F and Supplemental Figure 6, E and F). As cells with CA divide the resulting daughter cells often display low levels of chromosome mis-segregation and aneuploidy, which activates the DNA damage response pathway (*60*). Indeed, untreated *Pkd1*^*RC/RC*^ mice display elevated levels of γ-H2AX (Figure 6, G and H and Supplemental Figure 6, G and H), a surrogate marker of the DNA damage response. In contrast, there was a significant reduction in γ-H2AX staining following treatment with CCB02 or PJ34. We interpret these data to suggest that inhibiting centrosome clustering causes elimination of cells with CA from the population via apoptosis.

**Figure 6.**
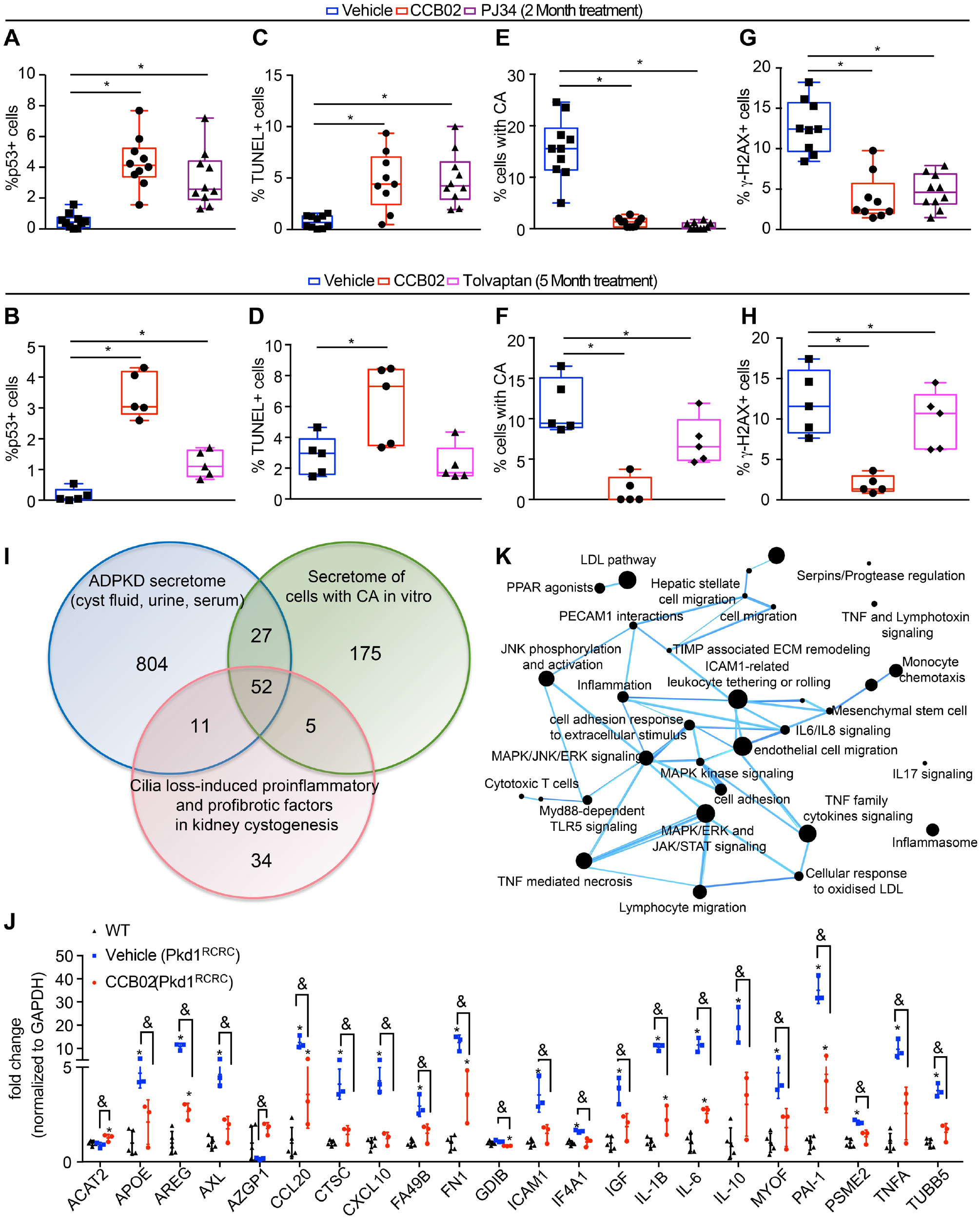
Centrosome clustering inhibitors promote p53-mediated apoptosis and attenuate pro-inflammatory signaling pathways. **(A** and **B**) Quantification of p53-positive nuclear staining in kidneys of *Pkd1*^*RC/RC*^ mice treated with CCB02 or PJ34 for 2 months or with CCB02 or Tolvaptan for 5 months. (**C** and **D)** Quantification of TUNEL staining in *Pkd1*^*RC/RC*^ kidney sections treated as indicated. (**E** and **F)** Quantification of the percentage of cyst-lining cells with excess (> 2) centrosomes. (**G** and **H)** Quantification of γ-H2AX-positive cells after treatment for 2 or 5 months. N = 10 mice per group (2-month treatment); N = 5 mice per group (5-month treatment). **(I)** Venn diagram showing the comparison of secreted factors identified in ADPKD patients and mouse models, secretome of cells with amplified centrosome, and cilia loss-induced pro-inflammatory and pro-fibrotic factors implicated in cystogenesis. **(J)** Quantification of the relative change in gene expression levels of selected factors in kidneys of wildtype or *Pkd1*^*RC/RC*^ mice treated with CCB02. N = 3 mice per group. * = p<0.05 (one-way ANOVA). **(K)** The network of significantly enriched biological themes defined by a CompBio pathway analysis tool, comparing the differentially expressed genes between control and CCB02-treated groups. The size of a sphere is proportional to the CompBio enrichment score of its theme.

Finally, we tested whether loss of cells with amplified centrosomes would attenuate their paracrine-mediated effects. Recent studies have shown that CA promotes the secretion of factors with proliferative, pro-inflammatory and pro-invasive properties (*14, 61-63*). This abnormal secretion is in part due to a stress response that results from increased reactive oxygen species (ROS) downstream of centrosome amplification (*14*). Therefore, the presence of amplified centrosomes can also influence the growth of adjacent cells with normal centrosome number in a non-cell-autonomous manner, via secretion of cytokines, growth factors and extracellular vesicles, suggesting a broader role for cells with CA. Importantly, numerous studies of ADPKD have demonstrated that cystic epithelial cells similarly display secretory phenotypes, resulting in elevated levels of cytokines and other growth factors. These promote a heightened and sustained immune response which contributes to cyst growth and interstitial fibrosis (*64, 65*). Thus, we hypothesized that elimination of cells with CA would help attenuate some pathways associated with such secreted factors.

To identify the centrosome amplification-associated factors that may be implicated in driving cyst formation in ADPKD, we compared the secreted factors identified in ADPKD patients and animal models (*65-78*) to those characterized by cells with CA *in vitro* (*14, 61*) (Figure 6I and Supplemental Excel sheet 1). We also included in the analysis proteins that are secreted upon defects in cilia formation in the kidney, as those have also been shown to promote an elevated immune response that exacerbates the cystic disease phenotype and renal fibrosis (*65, 79-82*). The *in-silico* analysis yielded a total of 52 overlapping proteins (Figure 6I). We then performed qRT-PCR analysis of those genes on kidneys isolated from wild-type and *Pkd1*^*RC/RC*^ mice treated with either vehicle or CCB02 (Figure 6J and Supplemental Figure 6I). As predicted, most of the genes (39/52, 75%) are upregulated in *Pkd1*^*RC/RC*^ mice compared to wild-type control (Figure 6J and Supplemental Figure 6I). Intriguingly, 21/52 (40.4%) show reduced levels following treatment with CCB02 (Figure 6J and Supplemental Figure 6I). Utilizing the Compbio platform (*83, 84*), we analyzed the network of these significantly enriched differentially expressed genes to determine the biological themes impacted between the control group and CCB02-treatment group (Figure 6K). The majority of these factors (e.g., TNF-α, IL-1*β*, IL-6, AREG, PAI1) are broadly involved in extracellular matrix remodeling upon renal injury by driving profibrotic and proinflammatory signaling. Some factors (e.g., ICAM1, CCL20, PSME2, AXL) can also enhance proliferative signaling pathways (via MAPK/ERK, JNK, and JAK/STAT signaling, among others) that leads to cyst growth stimulation. Moreover, other elevated factors in our dataset are known to regulate immune cell activity and infiltration (e.g. GDIB, CTSC), which is also known to exacerbate and promote cyst growth (*82, 85-88*). In sum, inhibiting centrosome clustering causes elimination of cells with CA via p53-mediated apoptosis which attenuates a subset of pro-fibrotic, proliferative and cystogenic signaling.

## Discussion

In this study, we tested whether centrosome clustering inhibitors can selectively target cells with amplified centrosomes in ADPKD models and attenuate the disease phenotypes. Centrosome amplification has been widely observed in human ADPKD specimens (*9, 10*), and we determined that this phenomenon is evident in almost half of the cysts. Importantly, analysis of spindle morphology showed that, as these cells divide, they cluster their excess centrosomes and form a pseudo-bipolar mitotic spindle (Figure 1). This robust centrosome clustering mechanism likely helps the cells avoid mitotic catastrophe, allowing them to proliferate and form cysts. The strong centrosome clustering effect was also found in human ADPKD cultured cells, as well as the *Pkd1*^*RC/RC*^ mouse model (Figure 4), which indicates that the survival mechanism of centrosome clustering is conserved in ADPKD, similar to what has been noted for cancer cells with CA (*15*).

We previously demonstrated that inducing CA in a developing wild-type mouse is sufficient to cause cyst formation, fibrosis, and a decline in renal filtration function (*17*). Here, we emphasize this point by conditionally driving centrosome amplification in the kidneys of the slow-onset *Pkd1*^*RC/RC*^ mouse model, at stages when cystogenesis is not yet evident. Indeed, the induction of CA caused accelerated cyst formation and growth, increased interstitial fibrosis and a decline in kidney function (Figure 2). These results highlight the additive effects of centrosome amplification in the context of ADPKD-inducing mutations. As mutations in ADPKD genes have been shown to disrupt centrosome biogenesis and result in cells with amplified centrosomes (*9, 10*), the formation of these ectopic centrosomes may therefore act in a feedback loop of sorts, further impacting kidney cystogenesis. This also establishes the hypothesis that eliminating the cells with CA may have beneficial outcomes.

Along these lines, we first tested the effects of different centrosome clustering inhibitors on cultured human ADPKD samples *in vitro*. Indeed, *PKD1*-null cells with CA treated with either CCB02 or PJ34 formed multipolar and disorganized spindles in mitosis (Figure 3). The ectopic centrosomes were present at most excess spindle poles and there was a concurrent decrease in the proportion of cells with bipolar spindles, indicating the treatment was effective. The centrosome de-clustering caused a mitotic delay, activation of the spindle assembly checkpoint and elimination of cells with CA from the population via apoptosis. Therefore, centrosome clustering inhibitors are effective in ADPKD cells and can arrest the proliferation of the cystic kidney cells with CA, consistent with what the inhibitors were shown to do in cancer cells and solid tumors ^45,^(*46-48*).

Next, we tested the effects of these inhibitors using the slow-onset *Pkd1*^*RC/RC*^ mouse model. Two separate treatment schemes were used in these mice: one starting at 6 months of age, the other at 9 months. This allowed us to examine the functional and morphological consequences of starting the treatment at different stages and severity of cystogenesis. Treatment with these inhibitors blocked the clustering of centrosomes in the cystic kidneys of *Pkd1*^*RC/RC*^ mice, indicating that they are active on kidney cells *in vivo*. In both cases, treatment of mice with either CCB02 or PJ34 resulted in a significant decrease in kidney size, cyst index, cyst number and fibrosis, while kidney filtration function was maintained (Figure 4 and 5). These data indicate that inhibiting centrosome clustering during the both the slow, progressive growth stage and the rapid cyst-expansion stage results in reduced cyst growth, improved kidney morphology and preserved renal function. These compounds were well tolerated by the animals, who showed better weight management, feeding behavior and urination that those treated with Tolvaptan (Figure 5). This is not surprising since Tolvaptan has been reported to cause changes in water retention, fluid secretion, urination and weight maintenance (*54, 55*). Importantly, treatment with the centrosome clustering inhibitors, followed by a washout period where no drug was administered for two months, showed that the observed improvement in kidney morphology and function were maintained well after the cells with CA are eliminated.

Finally, we sought to determine whether loss of cells with amplified centrosomes would attenuate their paracrine mediated-signaling effects. Several studies have shown that centrosome amplification promotes the secretion of signaling factors with proliferative, pro-inflammatory and pro-invasive properties (*14, 61-63*). The presence of amplified centrosomes in a cell can influence the growth of adjacent cells with normal centrosome number in a non-cell-autonomous manner, via secretion of cytokines, growth factors and extracellular vesicles. Moreover, numerous studies of ADPKD have shown that cystic epithelial cells display secretory phenotypes, resulting in elevated levels of cytokines and other growth factors. Indeed, recent studies have noted that a heightened and sustained immune response plays a key role in cyst growth and interstitial fibrosis (*64, 65*). *In silico* analysis of secreted factors in ADPKD and cancer cells with CA identified 52 common secreted factors, some of which we hypothesized may be impacted by elimination of cells with CA. qRT-PCR analysis indicated that expression of nearly half of those genes is reduced in *Pkd1*^*RC/RC*^ mice treated with a centrosome clustering inhibitor (Figure 6). The majority of these factors have been previously shown to be involved in extracellular matrix remodeling upon renal injury, by modulating profibrotic and proinflammatory signaling (*65, 79-82*). Some secreted factors are known to enhance proliferative signaling pathways that lead to cyst growth stimulation. Moreover, other elevated factors in our dataset are known to regulate immune cell activity and infiltration, which is known to exacerbate and promote cyst growth (*82, 85-88*). Thus, our qRT-PCR analyses helped to shed light on a subset of the centrosome amplification-dependent paracrine signaling pathways that are active in ADPKD.

In sum, our results indicate that inhibiting centrosome clustering in slow-onset ADPKD mice can block the proliferation of cells with excess centrosomes, results in reduced cyst growth, improved kidney morphology and preserved renal function. These data pave the way for testing of such compounds in patients with ADPKD.

## Author contributions

TC and MM conceived and designed the study. TC performed the vast majority of experiments presented in this study. AM and JG conceived of, and performed, the cell culture experiments. EL performed the qRT-PCR experiments and data analysis. KS assisted with mouse crosses and genotyping. TC and MM wrote and edited the manuscript. All authors reviewed the final version of the manuscript.

## Acknowledgments

We thank Dr. Sanjay Jain and the Kidney Translational Research Center (Washington University, St. Louis, MO) for help in procuring ADPKD patient specimens. We thank Dr. Renata Basto (Institut Curie, Paris) for generously sharing the mCherry-Plk4 mouse model. We also thank members of the Mahjoub lab and the Washington University Cilia Group for feedback on this project, and critical reading of the manuscript. Finally, we thank Dr. J. Fitzpatrick and the Washington University Center for Cellular Imaging (WUCCI) for assistance with some image acquisition, supported in part by Washington University School of Medicine, the Children’s Discovery Institute of Washington University and St. Louis Children’s Hospital (CDI-CORE-2015-505 and CDI-CORE-2019-813), and the Foundation for Barnes-Jewish Hospital (3770 and 4642). This study was supported by funding the Department of Defense PRMRP (W81XWH-20-1-0198) to M.R.M. The authors declare no competing financial interests.

## Materials and Methods

### Analysis of human ADPKD specimens

Human kidney samples were obtained from archived materials stored in the Kidney Translational Research Center (Washington University in St. Louis) according to the guidelines of a protocol approved by the Institutional Review Board of the Washington University School of Medicine. Renal specimens were fixed in formalin and embedded in paraffin, and 10 µm sections were cut and placed onto microscope slides. Sections from eight ADPKD patients, with ages ranging from 56–75 yr, were used for analysis. Specimens were processed and stained with antibodies to mark centrosomes and mitotic spindles as described below.

### Drug administration

All animal studies were performed following protocols that are compliant with guidelines of the Institutional Animal Care and Use Committee at Washington University and the National Institutes of Health. The Pkd1p.R3269C knock-in mice (equivalent to human Pkd1p.R3277C mutation, named *Pkd1*^*RC/RC*^ hereafter) were originally generated in C57BL6/N background and were maintained in a heterozygous state (RC/+) by mating with C57BL6/N WT mice. Genotyping was performed by extracting tail DNA followed by PCR using the following primers (all three primer pairs share the same forward primer): Forward: 5′-GTC TGG GTG ATA ACT GGT GT-3′; PKD1 full length (710 bp): Reverse: 5′-GGA CAG CCA AAT AGA CAG G-3′; PKD1-WT (480 bp): Reverse: 5′-AGG TAA CCC TCT GGA CTC T-3′; PKD1-Mutant (480 bp): Reverse: 5′-AGG TAA CCC TCT GGA CGC A-3′.

To test the effect of CCB02 and PJ34 on cyst progression at rapid stages of cystogenesis, 9-month-old *Pkd1*^*RC/RC*^ homozygous mice were administered either CCB02 (25 mg/kg, dissolved in 10 % PEG, 10% citric acid with a final DMSO concentration of 1.5 % [v/v]), PJ34 (10 mg/kg, dissolved in PBS) or solvent only (control) via gavage every other day for 2 months. Kidneys from 5 female mice and 5 male mice per group were harvested and analyzed at 11 months of age. To test the effects of CCB02 and Tolvaptan on cyst progression at early stages of cystogenesis, 6-month-old *Pkd1*^*RC/RC*^ homozygous mice were divided into three groups (control, Tolvaptan, CCB02). The mice were administered either CCB02 (25 mg/kg), Tolvaptan (10 mg/kg, dissolved in PBS) or solvent only (control) every other day via gavage for 5 months. Kidneys from 5 mice per group were harvested and analyzed at 11 months of age. To test the long-term benefits of treatment with the centrosome clustering inhibitors, nine-month-old *Pkd1*^*RC/RC*^ mice were administered CCB02 (25 mg/kg) every other day for 60 days, the treatment was stopped, the mice were followed for an additional 3 months until they reached 14 months of age and their kidneys (N = 4) harvested and analyzed.

### Histology, Immunohistochemistry, and Immunofluorescence

Both kidneys were isolated from each adult mouse as previously described (*17*) and fixed in one of two ways, depending on the downstream application. Specimens were fixed in 4% paraformaldehyde in PBS for 24 h at 4 °C, then embedded in paraffin. Samples were cut into 10 µm sections using a microtome (RM2125 RTS; Leica), placed onto microscope slides (Thermo Fisher Scientific), and stored at room temperature. For spindle morphology analysis, samples were cut into 15 µm sections. Alternatively, freshly isolated kidneys were placed in appropriately sized cryomolds (Tissue-Tek), immersed in Optimal Cutting Temperature compound (Tissue-Tek), and placed in a dry ice/ethanol bath for 10 min. Frozen samples were cut into 10 µm sections using a cryostat (CM1850; Leica), placed onto microscope slides, and stored at -80 °C. Histological staining of paraffin-embedded sections with hematoxylin and eosin (H&E) was performed using standard protocols.

For immunohistochemistry of paraffin-embedded sections, antigen unmasking was performed by boiling the slides in antigen-retrieval buffer (10 mM Tris Base, 1 mM EDTA, and 0.05% Tween-20, pH 9.0) for 30 min. For immunostaining of cryopreserved sections, samples were fixed using precooled (−20 °C) methanol for 10 min. After fixation, both types of fixed samples were pre-extracted with 0.05% Triton X-100 in PBS for 10 min at room temperature, incubated in blocking buffer (3.0% BSA and 0.05% Triton X-100 in PBS) for 1 h, followed by staining with primary and secondary antibodies. The complete list of antibodies used in this study is provided in Supplemental Table 1. All Alexa Fluor dye–conjugated secondary antibodies were obtained from Thermo Fisher Scientific and utilized at a final dilution of 1:500. Nuclei were stained with DAPI, and specimens mounted onto slides using Mowiol antifade mounting medium containing n-propyl gallate (Sigma-Aldrich). Images were captured using a Nikon Eclipse Ti-E inverted confocal microscope equipped with 10× Plan Fluor (0.30 NA), 20× Plan Apo air (0.75 NA), 60× Plan Fluor oil immersion (1.4 NA), or 100× Plan Fluor oil immersion (1.45 NA) objectives (Nikon). A series of digital optical sections (z-stacks) were captured using Hamamatsu ORCA-Fusion Digital CMOS camera at room temperature, and three-dimensional image reconstructions were produced. Images were processed and analyzed using Elements AR 5.21 (Nikon) and Photoshop software (Adobe).

### Evaluation of the number of centrosomes and mitotic spindle morphology

HK-2 and WT9-12 cells were grown on glass coverslips in multi-well plates in DMEM F-12 medium supplemented with 10% FBS. After treating the cells with CCB02, PJ34 or AZ82, cells were washed with PBS and fixed with ice cold methanol at -20 °C for 5 min. Fixed cells were blocked with 0.5% fish gelatin in PBS (block and wash buffer) for 60 min at room temperature or at 4 °C overnight, and then incubated with primary antibodies (centrosomal markers and spindle assembly checkpoint markers, as indicated in the figure legends) for 60 min at room temperature or at 4 °C overnight. After incubation, cells were washed with buffer three times for 5 min and incubated with species-specific Alexa Fluor-conjugated secondary antibodies (Thermo Scientific) at 1:1000 dilution for 60 min at room temperature. DAPI (1:1000; Invitrogen) was used to stain DNA. The following primary antibodies were used: rabbit anti Bub1 (1:50, #GTX107497), mouse anti γ-tubulin (1:500, Sigma # T6557), rat anti α-tubulin (*89*), mouse anti CPAP (*89*) and rabbit anti Cep152 (*89*). Images were collected with Leica SP8 laser scanning confocal microscope. Images were processed using Fiji/ImageJ, Adobe Photoshop and Adobe illustrator.

For quantification of centrosome number in kidney samples a minimum of five tissue sections—a midsagittal section and sections generated in 20 µm increments in both directions from the midsagittal section—were analyzed. Samples were stained with antibodies against γ-tubulin. Normal centrosome number was defined as cells containing one or two foci of γ-tubulin, and CA was characterized by the presence of more than two foci per cell. For evaluation of mitotic spindle morphology, paraffin-embedded kidney specimens were immunostained for γ-tubulin (centrosomes), α-tubulin (microtubules), pHH3 (mitotic marker), and DNA (DAPI). Three-dimensional reconstructions of z-stack images were used for the analysis. To determine the fraction of cells with abnormal spindle morphology, we counted all metaphase cells in each kidney section containing more than one centrosome and compared that with mitotic cells containing the normal complement of two centrosomes.

### Evaluation of Kidney Fibrosis

Masson’s trichrome stain and immunofluorescence staining using α-SMA were used for the assessment of fibrosis. Calculation of the total kidney area and the percentage area covered by α-SMA staining was utilized to measure the area covered by fibrotic cells. Multiple fields from each kidney section were imaged using a 20× objective. 15 randomly selected fields from three tissue sections per sample were quantified. The data are expressed as the mean area of α-SMA-positive foci per unit area (squared millimeter).

### Quantification of cyst number and cyst index

Sagittal kidney sections were stained with hematoxylin and eosin (H&E) and examined by light microscopy. ImageJ software was used to measure the area of each cyst and to quantify the total numbers of cysts. A dilated tubule was counted as a cyst if its area exceeded 0.02% of the total kidney area. Cysts were scored in three sections from each kidney sample.

### BUN and serum creatinine measurements

Blood serum was obtained by centrifugation (6,000 x g, 15 minutes at 4 °C) of blood samples isolated via sub-mandibular bleeding, and used for the BUN assay (BUN-Urea, BioAssay Systems) following the manufacturer’s protocol. Creatinine levels were quantified by HPLC (UAB-UCSD O’Brien Center, The University of Alabama at Birmingham).

### Analysis of DNA damage and apoptosis

Antibodies against γ-H2AX were used to identify DNA damage, and p53 staining was performed as described above. Images were captured using a Nikon Eclipse Ti-E inverted confocal microscope equipped with 40× Plan Fluor oil immersion (1.4 NA). The percentage of γ-H2AX-positive cells or nuclear p53-positive cells in each kidney section were determined. Briefly, ImageJ software was used to count the number of total cells (based on the DAPI blue stain) and the number of γ-H2AX-positive cells or nuclear p53-positive cells (based on the green stain) in each field. The number of green cells was then divided by the number of total cells for each field. The TUNEL method was used to quantify apoptosis following the manufacturer’s instructions (In Situ Cell Death Detection kit; Roche). The percentage of TUNEL-positive cells in each kidney was quantified as described above. All calculations were done from a minimum of five sections from experimental and control animals.

### RNA isolation, RT-PCR and quantitative PCR

Total RNA was isolated from mouse kidneys using Direct-zol™ RNA MiniPrep Plus (Zymo Research). 2 µg of RNA was reverse-transcribed using a High Capacity cDNA Reverse Transcription Kit (Applied Biosystems, ThermoFisher Scientific) according to the manufacturer protocol. Real-time PCR was performed with SYBR Select Master Mix (Applied Biosystems) in a 96-or 384-well plate format. 50 ng of cDNA was used in 10 ul final volume and the reactions run at the standard cycling mode recommended by the manufacturer, using QuantStudio 6 Flex system (Applied Biosystems). GAPDH was used as an endogenous control and data analyzed with ΔΔCt method. The complete list of primers used in this study is provided in Supplemental Excel sheet 2.

### CompBio analyses

The secreted factors identified by PCR were further analyzed using the CompBio (COmprehensive Multi-omics Platform for Biological InterpretatiOn) software package. CompBio analyzes single or multi-omic data entities (genes, proteins, microbes, metabolites, miRNA) and deliver a holistic, contextual map of the core biological concepts and themes associated with those entities. Conditional probability analysis calculated the statistical enrichment of biological concepts (processes/pathways) over those that occur by random sampling. Related concepts built from the list of differentially expressed entities were further clustered into higher-level themes (e.g., biological pathways/processes, cell types and structures, etc.).

### Statistical analyses

Statistical analyses were performed using GraphPad PRISM 7.0 or Microsoft Excel. The vertical segments in box plots show the first quartile, median, and third quartile. The whiskers on both ends represent the maximum and minimum values for each dataset analyzed. Collected data were examined by ANOVA or two-tail unpaired t-test as specified in the figure legends. Data distribution was assumed to be normal, but this was not formally tested. Statistical significance was set at P < 0.05.

---

**Supplemental Figure 1.**
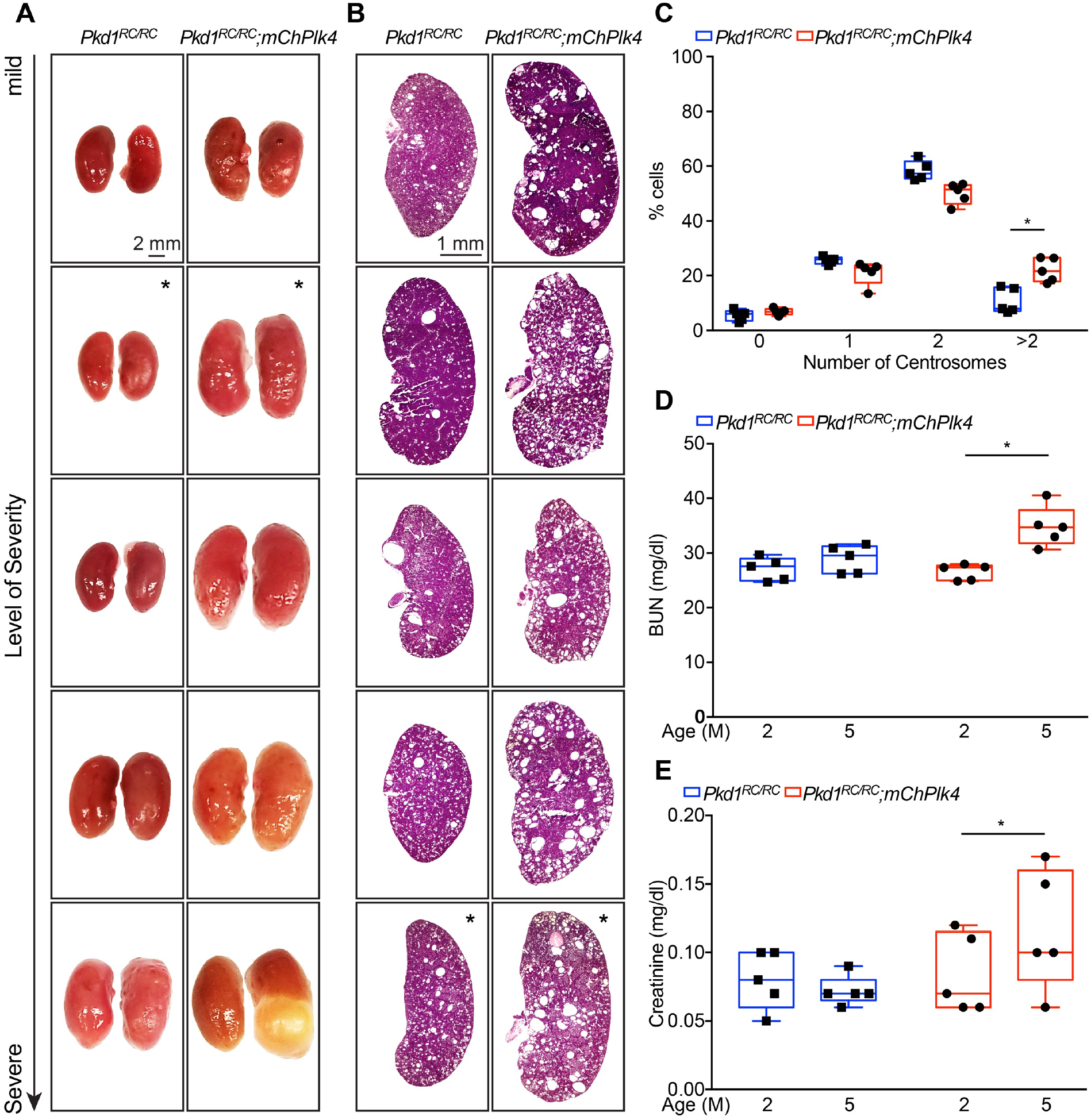
**(A** and **B)** Images of whole kidneys and H&E-stained sections of all *Pkd1*^*RC/RC*^ and *Pkd1*^*RC/RC*^*;mChPlk4* mice at 5 months of age that were analyzed in this study, organized by level of disease severity. Images containing an asterisk were the examples utilized in the main figure. (**C)** Quantification of centrosome number in cyst-lining cells of *Pkd1*^*RC/RC*^ and *Pkd1*^*RC/RC*^*;mChPlk4* kidneys. (**D)** Analysis of absolute levels of blood urea nitrogen (BUN) and (**E)** serum creatinine levels of *Pkd1*^*RC/RC*^ and *Pkd1*^*RC/RC*^*;mChPlk4* mice at 5 months. N = 5 mice per group for all experiments. * = p<0.05 (one-way ANOVA).

**Supplemental Figure 2.**
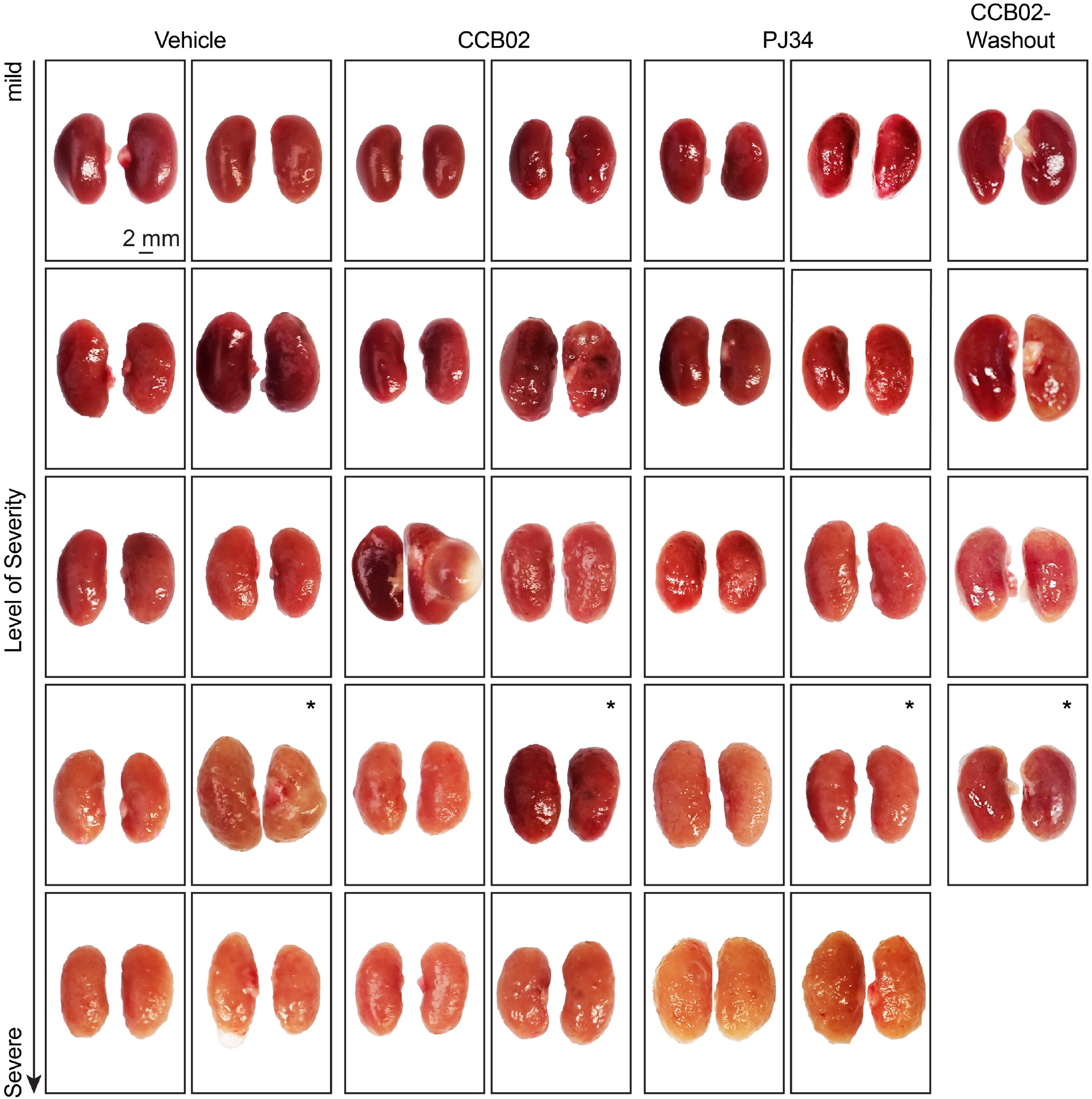
Images of whole kidneys from all *Pkd1*^*RC/RC*^ mice treated with either CCB02, PJ34, and post-washout that were analyzed in this study organized by level of disease severity. Images containing an asterisk were the examples utilized in the main figure.

**Supplemental Figure 3.**
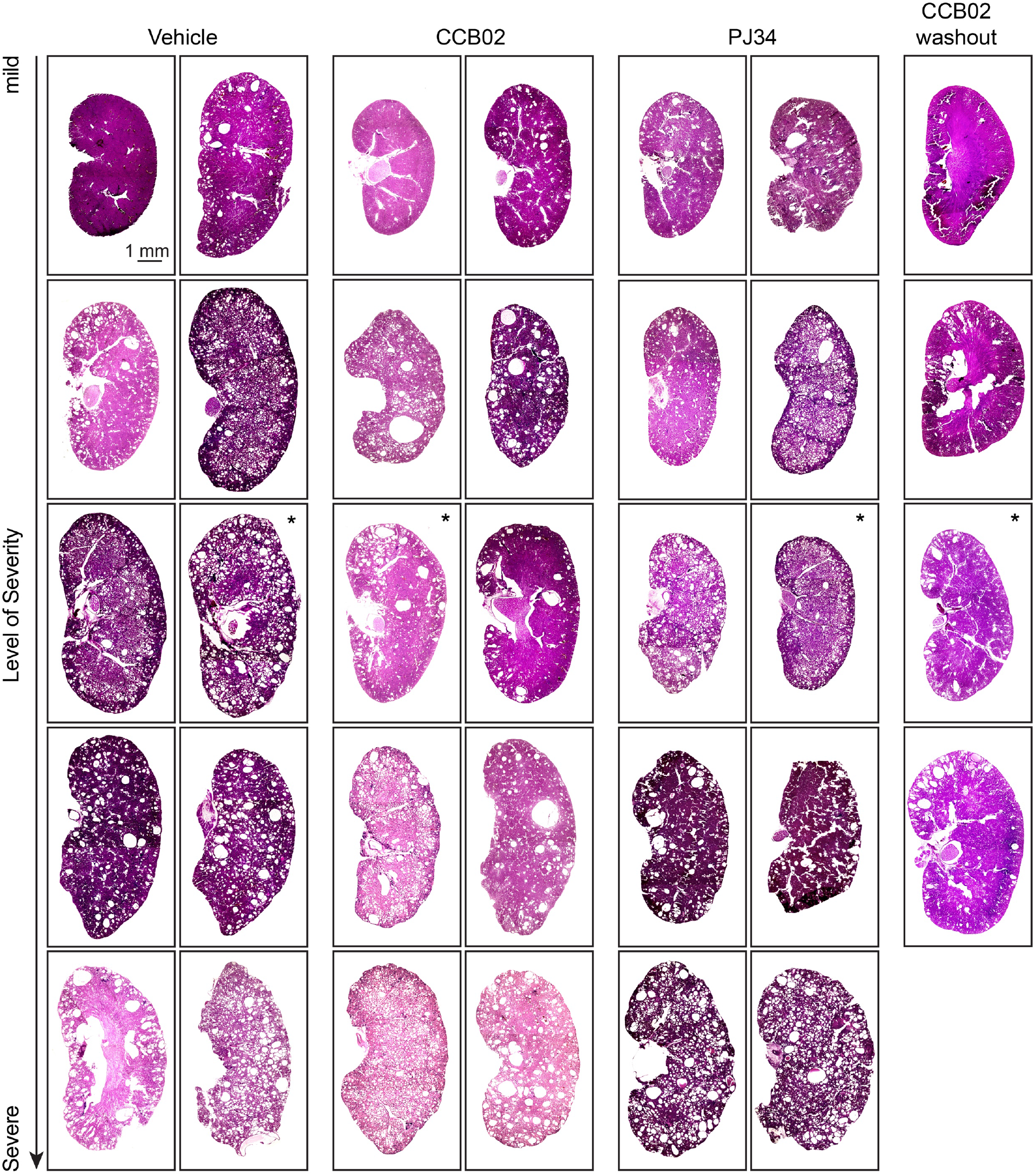
H&E stained images of kidney sections from all *Pkd1*^*RC/RC*^ mice treated with either CCB02, PJ34, and post-washout that were analyzed in this study organized by level of disease severity. Images containing an asterisk were the examples utilized in the main figure.

**Supplemental Figure 4.**
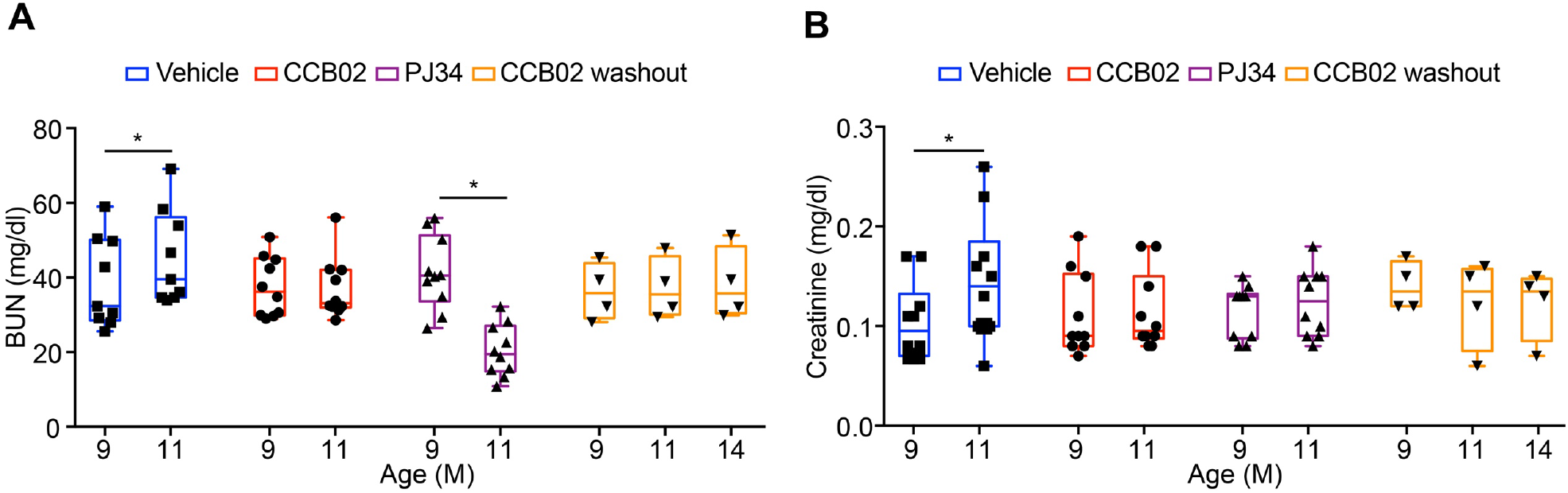
**(A)** Quantification of absolute levels of blood urea nitrogen (BUN) and (**B)** serum creatinine levels of *Pkd1*^*RC/RC*^ mice treated with CCB02 or PJ34 at 11 months, and post-washout (14 months). N = 10 mice each for Vehicle, CCB02 and PJ34; N = 4 animals for CCB02 washout group. * = p<0.05 (one-way ANOVA).

**Supplemental Figure 5.**
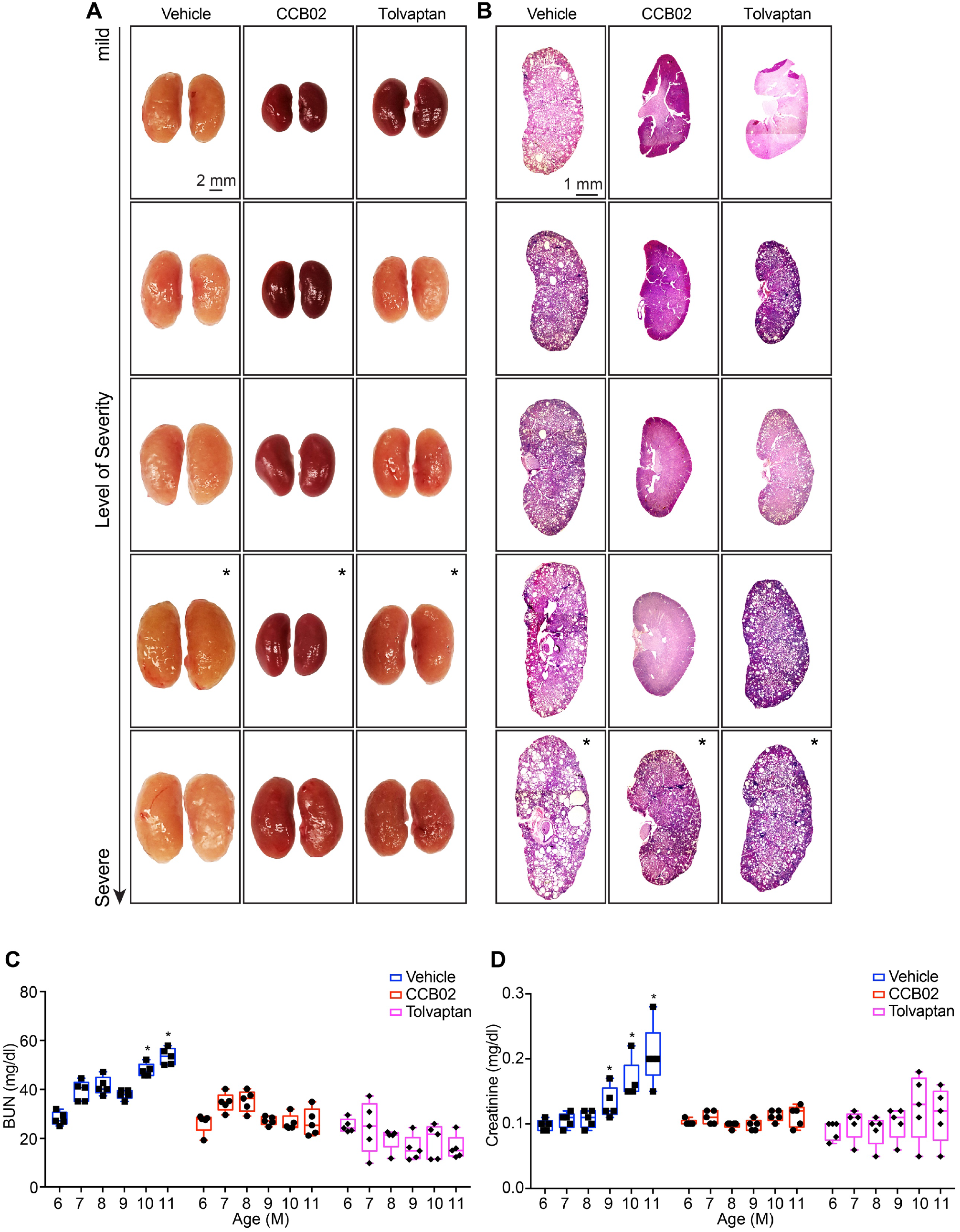
**(A)** Images of whole kidneys and (**B)** H&E-stained sections from all *Pkd1*^*RC/RC*^ mice treated with CCB02 or Tolvaptan that were analyzed in this study, organized by level of disease severity. Images containing an asterisk were the examples shown in the main figure. (**C** and **D**) Quantification of absolute levels of blood urea nitrogen (BUN) and serum creatinine levels of *Pkd1*^*RC/RC*^ mice during treatment timeline. N = 5 mice per group for all experiments. * = p<0.05 (one-way ANOVA).

**Supplemental Figure 6.**
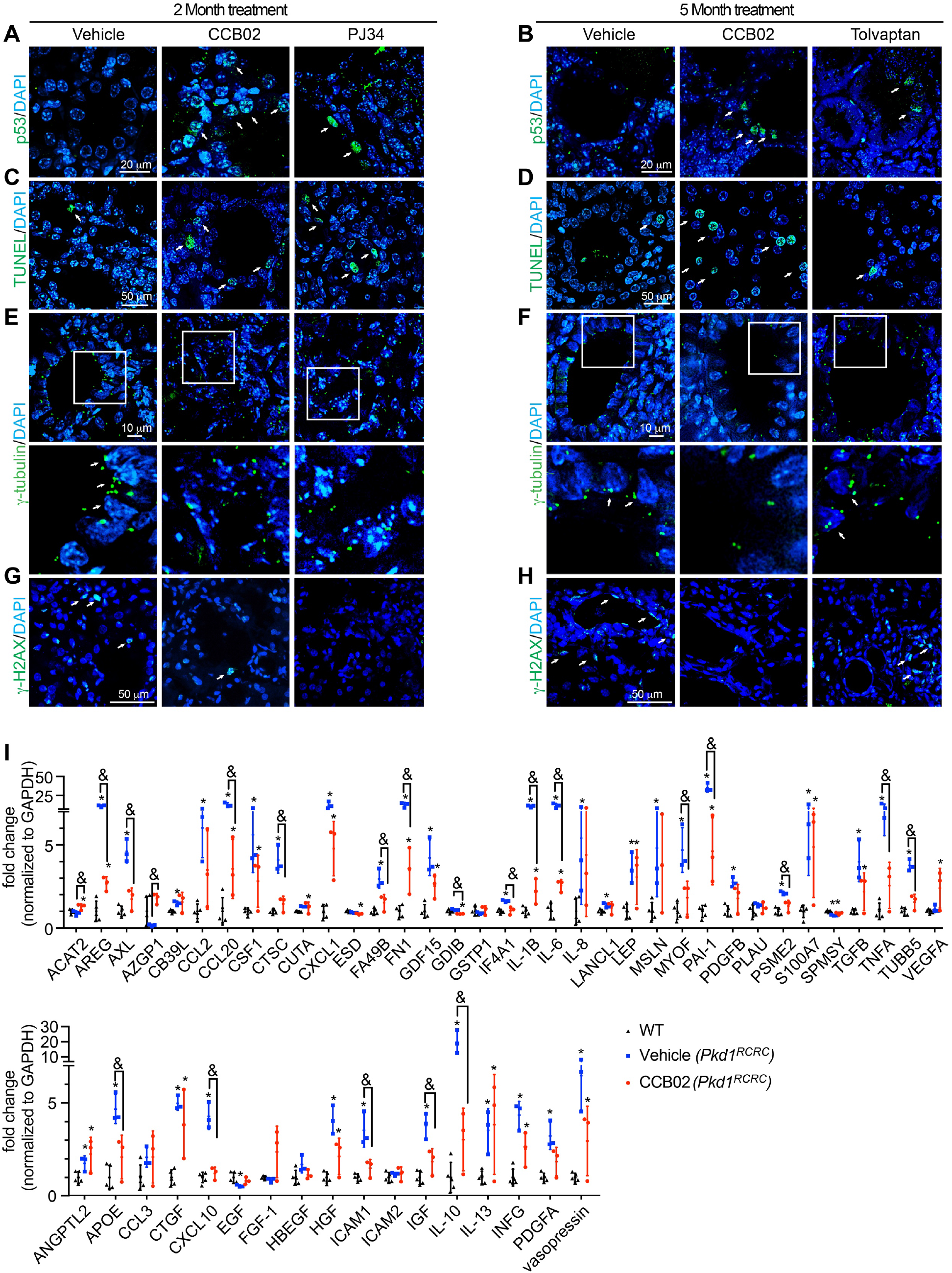
Immunofluorescence staining of kidney sections from *Pkd1*^*RC/RC*^ mice treated for either CCB02 or PJ34 for 2 months, or with CCB02 or Tolvaptan for 5 months. Kidney sections from 11 month-old mice were stained with antibodies to highlight nuclear p53 (**A** and **B**), TUNEL-positive cells (**C** and **D**), centrosomes (**E** and **F**), γ-H2AX (**G** and **H**), and DNA (DAPI). Arrows point to examples of positive cells in each case. White box in panels **E** and **F** highlight magnified regions shown below. (**I)** qPCR-based quantification of the relative change in gene expression levels of all 52 factors identified from *in silico* analysis. N = 3 mice per group. * = p<0.05 (one-way ANOVA).

**Supplemental Table 1:**
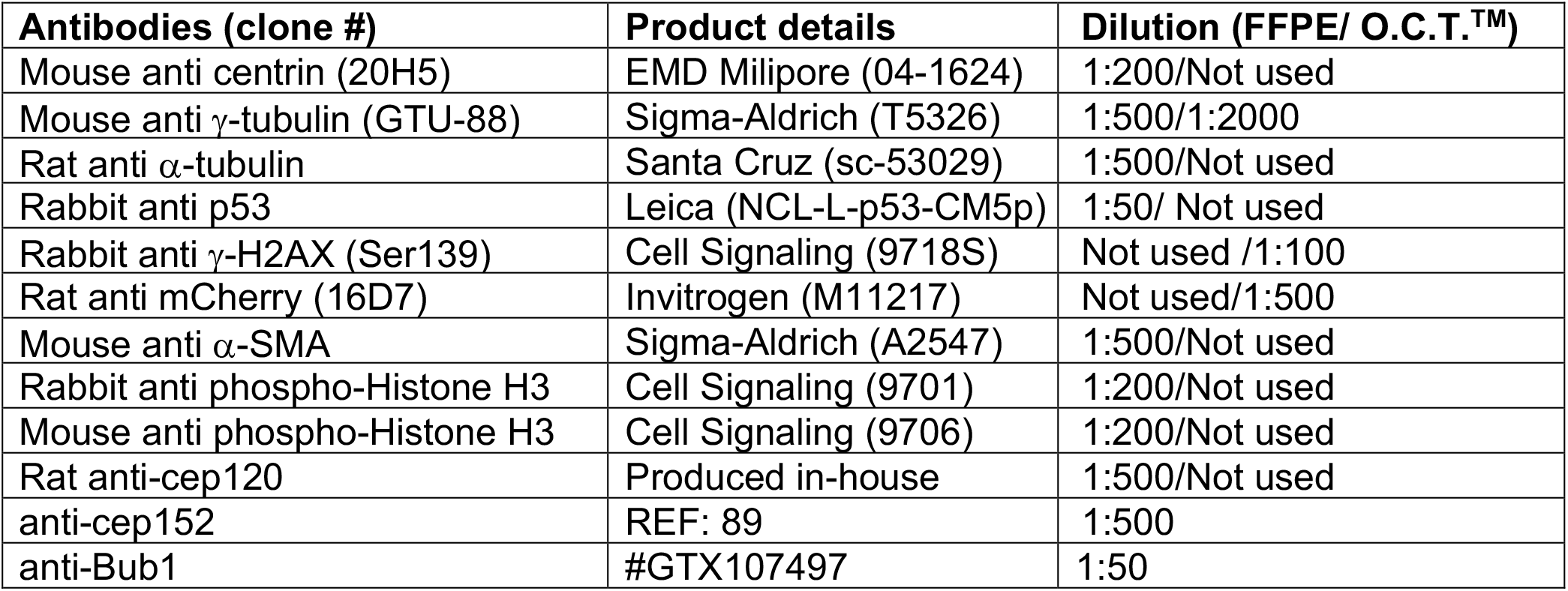
List of antibodies used in this study

| Antibodies (clone #) | Product details | Dilution (FFPE/ O.C.T.<sup>TM</sup>) |
| --- | --- | --- |
| Mouse anti centrin (20H5) | EMD Milipore (04-1624) | 1:200/Not used |
| Mouse anti γ-tubulin (GTU-88) | Sigma-Aldrich (T5326) | 1:500/1:2000 |
| Rat anti α-tubulin | Santa Cruz (sc-53029) | 1:500/Not used |
| Rabbit anti p53 | Leica (NCL-L-p53-CM5p) | 1:50/ Not used |
| Rabbit anti γ-H2AX (Ser139) | Cell Signaling (9718S) | Not used /1:100 |
| Rat anti mCherry (16D7) | Invitrogen (M11217) | Not used/1:500 |
| Mouse anti α-SMA | Sigma-Aldrich (A2547) | 1:500/Not used |
| Rabbit anti phospho-Histone H3 | Cell Signaling (9701) | 1:200/Not used |
| Mouse anti phospho-Histone H3 | Cell Signaling (9706) | 1:200/Not used |
| Rat anti-cep120 | Produced in-house | 1:500/Not used |
| anti-cep152 | REF: 89 | 1:500 |
| anti-Bub1 | #GTX107497 | 1:50 |

**Supplemental Excel sheet 1**: Full list and comparison of secreted factors identified in various studies, data used in Figure 6 I-J.

**Supplemental Excel sheet 2**: Primer list for qRT-PCR of secreted factors.

